# Factors influencing birch (*Betula spp.*) age distribution in Norway: Insights from tree ring data

**DOI:** 10.1101/2025.06.10.658859

**Authors:** M Vergarechea, C Antón-Fernández, J.U Jepsen, O. P.L Vindstad, R Astrup

## Abstract

The age-class distribution of forests is a key indicator of both carbon stock potential and biodiversity conservation, playing a vital role in sustainable forest management. In Norway, birch (*Betula* spp.) is the most abundant tree species, covering 42% of the forest area. Understanding the factors that shape the age structure of birch is essential for developing management practices that balance timber production, carbon sequestration, and biodiversity conservation. Using data from the Norwegian National Forest Inventory (NFI), we examined the age-class distribution of birch trees across various site conditions. Our analysis revealed that middle-aged trees (50–100 years) were prevalent in most regions, while older trees were notably scarce, particularly in highly productive areas. This pattern reflects management strategies prioritizing younger, fast-growing trees to maximize economic returns. In contrast, less productive sites, which are often managed less intensively, tend to support older trees. Additionally, younger birch trees revealed significantly greater radial growth than older generations when evaluated at the same biological age (e.g., at 10 years old), particularly under favorable site conditions. These findings underscore the combined effects of site productivity, forest management, and environmental factors on growth dynamics and age-class distribution.

## Introduction

Reliable assessment of the age distribution of forests is essential for both understanding their historical context and anticipating their potential future development (Vilén et al. 2012). The age distribution of trees offers insight into different key aspects: it facilitates the estimation of potential harvesting yields (Verkerk et al. 2011), carbon stocks (Böttcher et al. 2008) and it may serve as an indicator of biodiversity, disturbance risk, or recreation value of forests (Liira et al. 2007, Schelhaas et al. 2010).

Tree age distribution is shaped by various factors, including seed availability and the type, severity, or frequency of natural disturbances (Kuuluvainen et al. 2002, Xing et al. 2012, Burch and Sánchez Meador 2018). Forest management practices also play a crucial role in shaping age structures, often resulting in stands with relatively uniform ages up to the rotation period—the point at which trees are considered mature for harvesting. (Fall et al. 2004, Wulder et al. 2009). In Fennoscandian countries, managed forests are typically harvested within economic rotation periods, which rarely exceed 80– 120 years for spruce and 50–60 years for birch (Wallenius 2002, Rytter et al. 2016a). These timeframes are designed to maximize timber yield and economic returns, preventing stands from reaching significantly older ages. As a result, only 13.5% of Finland’s forests and 11% of Sweden’s forests contain trees older than 120 years, primarily due to clear-cutting practices that remove trees long before they reach their maximum biological lifespan (Skogsstatistisk 1998).

Quantifying tree age can be challenging since it cannot be directly observed in the field. Instead, it is estimated by analyzing annual growth rings obtained with an increment borer (Baker et al. 2005, Dang et al. 2010). Most research using tree ring data has historically been focused on reconstructing past climates or examining tree species responses to climate and disturbances (Gomes Marques et al. 2018, Sanchez-Morales et al. 2019, Altman 2020). Furthermore, in most dendroecological and growth-trend studies, data are collected from a selection of sites, which may not accurately represent the entire range of variability within the population. In contrast, National Forest Inventories (NFIs) can provide accurate information on the main dendrometric variables, including age (Breidenbach et al., 2020). The establishment of NFIs started in the Nordic countries during the late 1910s and early 1920s, with additional European countries developing their inventories after World War II (Tomppo et al. 2010, Vilén et al. 2012). Several European NFIs had integrated increment core sampling into their methodologies by that time. Today, the use of this technique is likely even more widespread, underscoring its role in improving the accuracy and reliability of forest data collection. The Norwegian NFI protocol (Breidenbach et al. 2020) specifies that for each permanent plot, an increment core should be extracted from one or more representative trees of the dominant species situated just outside the plot. These tree species include Norway spruce (*Picea abies* (L.) Karst), Scots pine (*Pinus sylvestris* L.) and birch (*Betula* ssp.).

Using NFI data covering all of Norway, we examined the age distribution of Silver birch (*Betula pendula* Roth.) and Downy birch (*Betula* p*ubescens* L.). Downy birch, which forms the alpine and arctic forest lines, dominates birch forests in Norway, comprising over 95% of the total birch volume (Smith et al. 2014). In contrast, Silver birch has a more southerly distribution and occurs at lower elevations. For simplicity, we collectively refer to both species as "birch" in this manuscript. Both species are pioneer trees well adapted to colonizing open areas. However, their distribution can be limited by several environmental factors such as air and soil temperature, or snow cover (Aas and Faarlund 2000, Wielgolaski et al. 2004, Holtmeier and Broll 2005). In most forest clear-cuts, birch seedlings naturally regenerate and establish spontaneously, contributing to the early successional dynamics of the ecosystem. Birch trees are well-adapted to oligotrophic soils but require abundant light for optimal growth (Bandekar and Odland 2017). Thus, as a light-demanding pioneer tree species, birch is characterized by a high youth growth and negatively affected by competition for light with neighboring trees (Vanhellemont et al. 2016). The typical lifespan of birch ranges from 50 to 100 years, although some individuals may exceed this age (Cameron 1996). In boreal forests, the species provides key ecosystem services, including timber and bioenergy production, as well as serving as an important fodder source for ungulate foraging (Lidman et al. 2024). In Scandinavia, it is common to find birch growing in mixtures with other species, particularly Norway spruce and Scots pine (<u>Johansson, 2003</u>).

This birch admixture enhances ecological sustainability while also offering forest owners economic and management benefits, as naturally regenerated broadleaved species in conifer-dominated plantations are a key component of sustainable forest management and a requirement of certification standards (FSC 2010, 2020). In alpine and subarctic birch forests, key disturbances of these birch forest include insect outbreaks, leading to mass mortality (Tenow 2005, Jepsen et al. 2013), avalanches and extreme weather events (Hansson et al. 2021). Additionally, anthropogenic disturbances, such as grazing by sheep and semi-domestic reindeer, significantly impact these forests, particularly in the subalpine and subarctic region (Nygaard et al. 2022).

Birch is of economic and ecological importance in the Nordic countries (Hynynen et al. 2009), where it constitutes between 11% and 16% of the total growing stock volume. However, in Norway, its use in timber production remains limited (Sjølie, 2011). Instead, birch is primarily utilized for fuelwood and bioenergy markets, driving the development of mechanized harvesting techniques tailored for processing small-dimension birch wood (Holmström 2015), typically under a 50–60-year rotation cycle (Cameron 1996, Rytter et al. 2016b). In Western Europe, the merits of birch are not yet generally appreciated by foresters. Towards its southern ranges, the species is valued more for its role in supporting biodiversity and enhancing soil conditions (Vanhellemont et al. 2016). The emphasis on Norway spruce and Scots pine in Norwegian forestry has significantly influenced the age and size distribution of birch forests, particularly in high-productivity areas (Sjølie 2011). However, the relatively limited management of birch forests provides a unique opportunity for detailed analysis of their age distribution and growth dynamics, in contrast to species subjected to more intensive forestry practices. Understanding stand age distribution is crucial not only for assessing current carbon sequestration potential but also for predicting future forest development and long-term ecological dynamics.

The tree ring dataset used in this study spans a latitudinal gradient from 58 to 71° N across five distinct vegetation zones, offering a unique opportunity to investigate the distribution of birch tree age across Norway. Comparable datasets are scarce, with Sweden being one of the few countries with similar data (Fridman et al. 2014). Therefore, the primary aim of this study is to analyze the factors that have influenced the current age structure of birch stands across all the major bioclimatic gradients in Norway. Additionally, we compare growth rates (radial growth) across age groups and under varying growing conditions, including site index and biogeographical regions. Understanding the variations in birch growth potential across different age groups is particularly valuable (Vanhellemont et al. 2016) as it could provide insights into how birch trees respond to environmental conditions and management practices. Specifically, we address the following research questions: (1) What is the current age-class distribution within birch forests across Norway? (2) What factors influence the distribution of birch ages across the different biogeographical regions? (3) To what extent do these factors drive differences in growth rates between younger and older birch trees in Norway?

## Materials and Methods

### Study region

The study covers the entire mainland of Norway, ranging from 58 to 71° N and from 5 to 31° E. In Norway, forests cover about 14 million ha, representing 37.5% of the land area, of which 8.3 million ha are productive forests with a mean growing stock of 105 m^3^ha^-1^ on average and mean growth of 2.6 m^3^ha^-1^yr^-1^ . Deciduous trees, particularly birch, dominate the forest area in Norway (42%), although the coniferous species Norway spruce and Scots pine make up the majority of biomass (42% and 30%, respectively) (SSB 2020).

Norway exhibits a high diversity in biogeographic and climatic conditions. In this study, we adopted the methodology of Bakkestuen et al. (2008) to categorize Norway’s landscape into five major vegetation zones (Fig. 1). These areas span a diverse range of ecosystems, from boreal to alpine. They are characterized by distinct species compositions, which serve as indicators of the varying precipitation and temperature conditions unique to each zone (Moen 1998). The boreal zone generally experiences a climate with cold winters, prolonged snow cover, and a short growing season. In Norway, this zone is divided into four subzones—boreonemoral, southern boreal, middle boreal, and northern boreal (Fig. 1)—based on varying climate gradients. The boreonemoral is affected by agricultural land-use, and the vegetation is fragmented, consisting of pastures and a mosaic of deciduous forest with, for example, *Quercus* spp L., *Acer platanoides* L., *Fraxinus excelsior* L., *Tilia cordata* Mill., and *Ulmus glabra* Huds., and boreal forests with *P.abies* and *P. sylvestris*. The southern-boreal zone, can be difficult to differentiate from the boreonemoral, since it is also affected by land-use and, as in the previous case, presents localized areas with warmth-demanding deciduous trees. In the southern- boreal zone, trees and shrubs such as *B. pubescens* and *Salix spp*. L. are more common than in the boreonemoral zone. The middle-boreal zone is mostly dominated by coniferous forest. However, deciduous trees, such as *B. pubescens*, *Alnus incana* L., *Sorbus aucuparia* L., and *Populus tremula* L., are also commonly found in the area. In contrast, warmth-demanding tree species are less frequent, occurring mainly in favorable locations such as south-facing slopes. In the transition from the northern- boreal zone to the alpine and arctic regions, the landscape is primarily covered by mountain-birch forests and conifers. The density of *P. abies* and *P. sylvestris* gradually decreases with rising elevation. Finally, the alpine region is predominantly characterized by a variety of shrubs, such as *Juniperus communis* L., *Betula nana* L. and *Salix spp.* along with dwarf-shrubs, graminoids and herbs.

**Figure 1.**
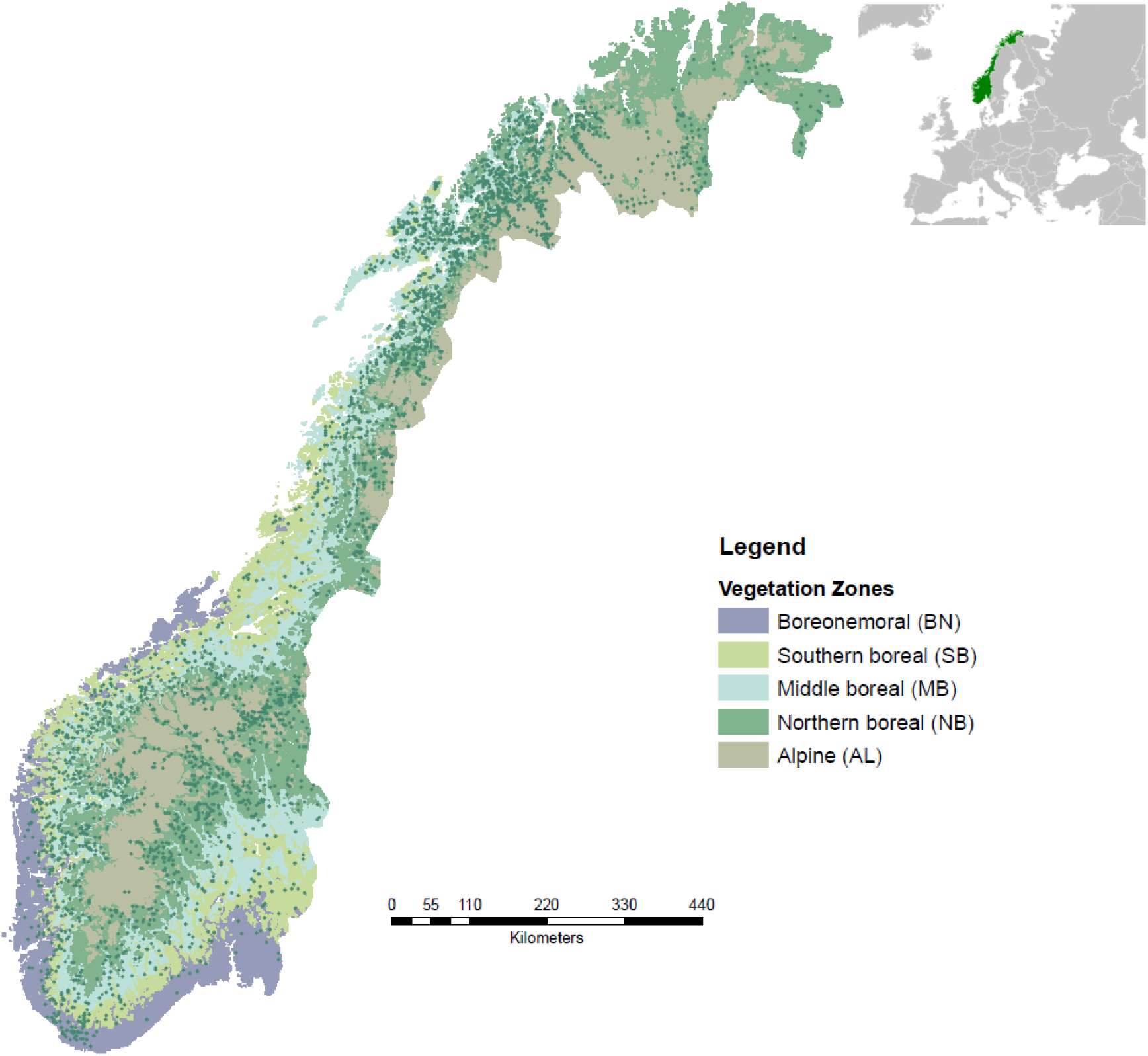
Vegetation zone map according to Bakkestuen et al. 2008. The NFI plots which contributed data to this study are shown as green dots.

### Field sampling and laboratory measurements under the National Forest Inventory

For the current study, we used data collected during 2005-2020 as part of the Norwegian NFI. The Norwegian NFI is based on permanent plots (Fig. 1) established at each intersection of a 3 x 3 km grid in the lowlands, and a 3 x 9 km grid in the mountains. In the northernmost region (Finnmark) a 9 x 9 km grid is used throughout (Breidenbach et al. 2020). The plots are circular, with a size of 250 m^2^ and resampled every 5^th^ year with 1/5 of all NFI plots visited annually. Within the 250 m^2^ plots, diameter at breast height (DBH), species, vitality status, and position are recorded for all trees with a DBH larger than 5 cm. Site index (SI) is included in NFI based on the H40system, which uses dominant tree height at a base age of 40 years at DBH as the SI value (for a more detailed description of the SI estimation, see Breidenbach et al. 2020). Thus, each NFI plot has a site index value (6, 8, 11, 14, 17, 20, 23 and 26) for the dominant species. In this study, we used the site index associated with each NFI plot, and we grouped the high SI plots in a group called “high SI” (SI > 17) due to the low representation of high values in our samples.

The age of each NFI plot was estimated using increment cores taken from one or more representative trees located just outside the survey plot. The representative trees were identified based on the dominant species in the plot, which was determined by the highest timber volume over bark (m^3^ ha^−1^) in each NFI plot, according to Breidenbach et al. (2020b). Consequently, we selected 2785 NFI plots where birch was recorded as the dominant species. Within these plots, increment cores were collected from a total of 2818 birch trees. The cores were air-dried, mounted onto wooden slats, and sanded to obtain a smooth cross-section. Tree ring widths (TRW) were measured using a LINTAB 6 tree-ring measurement station and TSAPWin version 4.81c software. Finally, the age of each tree at breast height was determined by counting the number of rings from the outermost ring to the pith. Visual comparison of ring-width graphs was used as a quality control procedure, however, after this procedure, 14 trees were removed for the sample due to inconsistences in tree ring patterns. The dataset may thus contain a certain amount of noise due to the undetected presence of missing or, less commonly, false rings, which may occur in birch (Cairns et al. 2012). However, we consider this of minor importance in the context of the current study, where the focus is on evaluating regional patterns in age distribution, rather than the identification of specific pointer years, or changes in growth patterns due to climate variations. This approach allowed us to preserve a large enough sample size to detect changes in the age distribution of birch forests.

### Analysis of age distribution and growth

To evaluate differences among age classes, each tree was assigned to its respective age group: *old* (age > 80 yrs), *intermediate* (35 yrs < age ≤ 80 yrs), or *young* (age ≤ 35 yrs). Therefore, using this age classification, we determined the distribution of trees across these three age classes within each vegetation zone in Norway. This was done both in terms of absolute numbers (by counting the samples) and as a proportion of the area covered by each age class. For the latter analysis, we assumed the age of the representative tree (the sampled tree) to reflect the age of the entire inventory plot. In addition, the NFI categorizes Norwegian forests into four different strata, each with varying sampling intensities as described by Breidenbach et al. (2020) (see section 2.2). As a result, the size of the forest area represented by each plot varies across these strata, with each plot corresponding to a specific portion of the forest. As the NFI represents a systematic sample with known inclusion probabilities, we can calculate the proportion of area occupied by each age class (old, intermediate and young) within each vegetation zone with the standard NFI estimators (Breidenbach et al. 2020).

We further examined how birch tree ages are distributed across various site index values and vegetation zones. For this purpose, we divided the ages into 10-year intervals, rather than the previous year age classes.

The TRW series were subsequently transformed into basal area increments (BAI) as this variable is more effective for capturing growth trends and accounting for variations due to the increase in tree size and age. BAI was calculated as in Eq. (1). Then, to facilitate comparisons of time-dependent growth characteristics, the TRW and BAI measurements were aligned according to biological age, ensuring that growth rings from different trees correspond to the same stage of development. To achieve this, the TRW and BAI data were grouped and averaged over site index, vegetation zone, and age class (old, intermediate, and young) using the arithmetic mean. Subsequently, the data were smoothed to emphasize the long-term growth trends. Specifically, we performed an average smoothing, where TRW and BAI data were averaged over fixed intervals of 5 years.

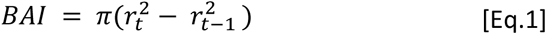

Where 𝑟_t_ and 𝑟_𝑡−1_ are the stem radius at the end and at the beginning of a given annual ring increment.

## Results

### Variation in tree age structure

There is a clear trend of increasing average age from the boreonemoral zone to the alpine zone (Table 1), moving from 50.60 years in boreonemoral zone to 79.75 years in alpine zone. This trend is also shared by the maximum age. The highest standard deviation in the middle boreal (MB) indicates that this zone has the greatest variability in ages.

**Table 1.** Descriptive summary of the age (breast height) distribution across vegetation (biogeographical) zones. Here BN = boreonemoral, SB = southern boreal, MB = middle boreal, NB = northern boreal and AL= Alpine.

|  | BN | SB | MB | NB | AL |
| --- | --- | --- | --- | --- | --- |
| **Number of samples** | 131 | 319 | 842 | 1243 | 269 |
| **Average age** | 50.69 | 61.72 | 73.74 | 76.55 | 79.75 |
| **sd** | 26.31 | 31.24 | 33.27 | 31.76 | 30.58 |
| **Maximum age** | 110 | 138 | 177 | 189 | 201 |
| **Minimum age** | 10 | 8 | 10 | 11 | 15 |

The analysis of tree age distribution across vegetation zones revealed distinct differences in age structure (Table S1). In Norway, the intermediate age class has the highest representation in all zones, with particularly large sample sizes in the northern boreal (NB) zone (766 trees) and the middle boreal (MB) zone (472 trees). The young age class is underrepresented in comparison, with the alpine (AL) zone having the fewest samples (21 samples). The old age class shows a higher presence in the middle boreal (402 samples) and northern boreal (540 samples) zones, while the northern boreal (28 samples) and alpine (130 samples) zones have much lower representation. Growth rates (mean ring width) consistently decline with age, with the alpine zone displaying the lowest growth rates in all age classes, likely due to harsher environmental conditions.

Histograms show a higher proportion of trees in the intermediate age class in northern boreal compared to other sites (Fig. 2). Conversely, vegetation zones such as boreonemoral and alpine exhibit broader age class distributions with a less pronounced peak and a more diverse age structure.

**Fig. 2.**
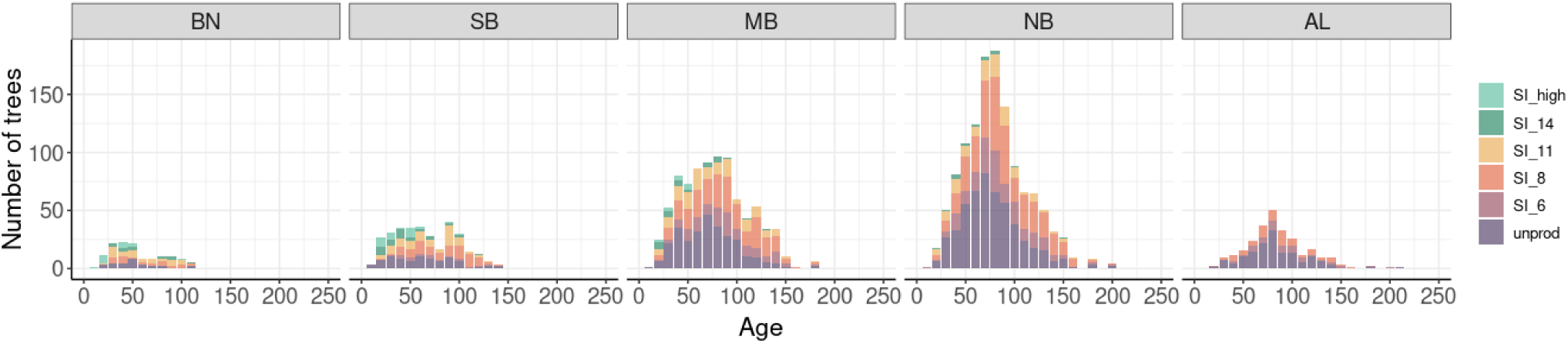
Histograms for the number of trees within each age class across the study sites, indicating the age structure within each site. Here BN = boreonemoral, B = southern boreal, MB = middle boreal, NB = northern boreal and AL= Alpine

The density plots illustrate a general trend where higher site indices show younger age structure (Fig. 3). Particularly, the SI_high and SI_14 groups display a notable peak in the younger age classes, and site index values above 11 were practically absent in age classes beyond 80-100 years (Fig. 2). Regarding the vegetation zones, the highest site index populations (SI high = 17, 20, 23 classifications) are found in the oceanic west coast areas, specifically boreonemoral, southern boreal, and middle boreal, although with a relatively small presence. In contrast, lower site indices, such as SI_6 and the nonproductive group, show a more uniform distribution across age classes with a slight increase in middle-aged trees.

**Figure 3:**
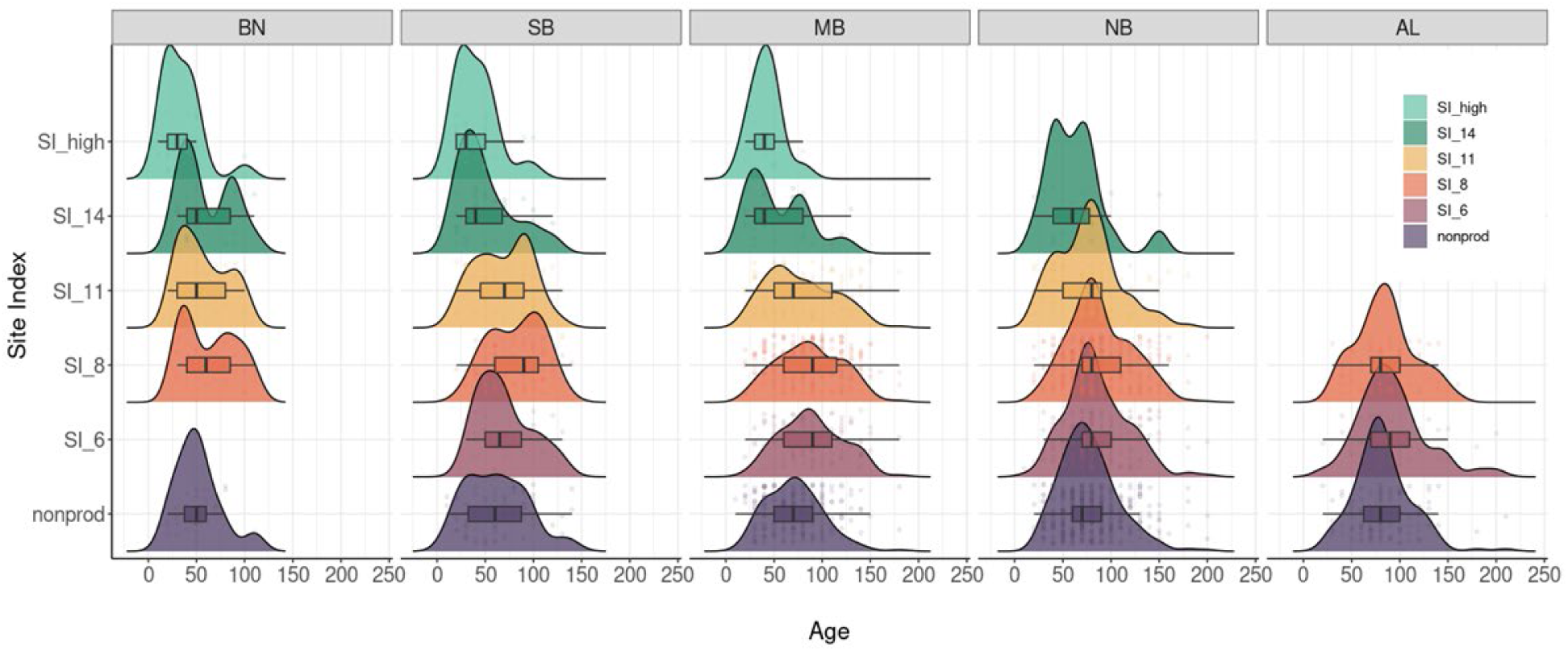
Age class distribution of birch tree populations across different site index (SI) groups, as well as across different vegetation zones. “nonprod” denotes nonproductive plots. BN = boreonemoral, SB = southern boreal, MB = middle boreal, NB = northern boreal and AL = Alpine

When examining radial growth across different age classes, we found that younger birch trees revealed significantly greater radial growth than older generations when evaluated at the same biological age, particularly under favorable site conditions. For example, a younger tree (currently 35 years old) grew faster at 10 years of age than an older tree (currently 80 years old) when both were at the same biological age of 10 years. As illustrated in Figure 4, in areas with a site index of 14, trees from the oldest age class achieved an average tree-ring width (TRW) of only 2 mm per year when they were 10 years old. In contrast, trees from the youngest age class attained an average TRW of 2.8 mm per year at the same biological age. In terms of basal area, this corresponds to an average of 3 cm² for older trees and 13.6 cm² for younger trees (Fig. S4). This highlights the enhanced growth potential of younger generations, particularly under favorable site conditions. Growth differences in both TRW and basal area between intermediate and older age classes were pronounced during the first twenty years in plots with a site index of 14 or higher. In contrast, in plots with a lower site index, both age classes exhibited similar growth patterns.

**Figure 4:**
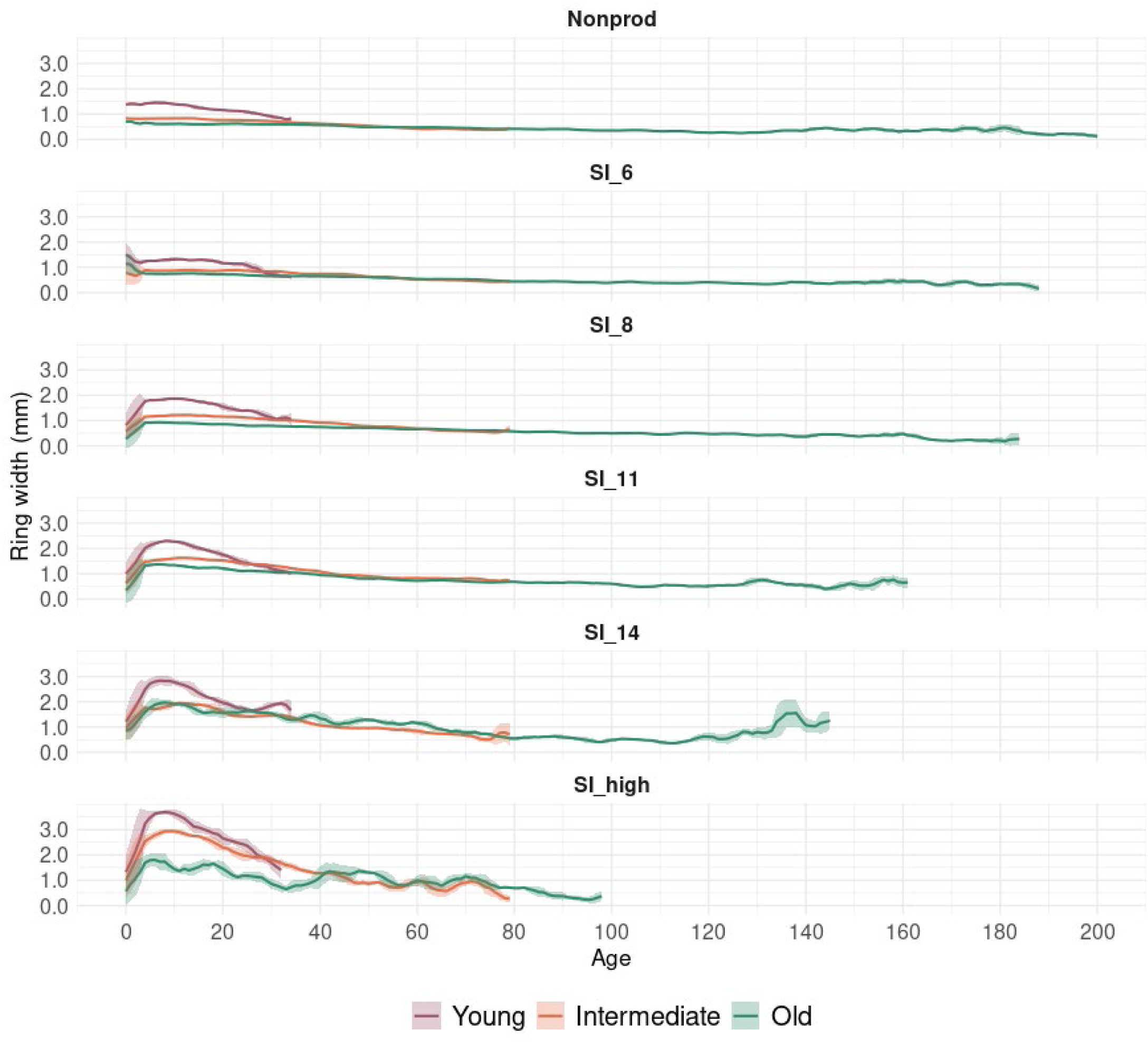
Growth trends for different site index for young (≤ 35 yrs), intermediate (35-80 yrs) and old age (> 80 yrs) classes based on cambial age. The shaded area represents the standard deviation of the average tree-ring widths.

Average growth trends of the different biogeographical zones show similar patterns over the full lifespan of the trees (Fig. S3). TRW initially increases, reaching a maximum after about 15 years, and then decreases until about the age of 100 years, where site sample replications drop, and growth curves become noisier. The greatest growth differences between zones occur at an average age of 10 to 15 years, with the widest tree rings (2.23 mm) observed in the boreonemoral zone and the narrowest rings (0.77 mm) in the alpine zone.

## Discussion

As highlighted in previous studies, accurately determining tree age is inherently challenging due to limitations such as the difficulty of direct measurement, and the potential costs involved. As a result, few studies have examined the age distribution of tree species across broad environmental gradients, such as those found in Norway. In our study, we observed significant variations in the age structure of birch across different biogeographical regions and site conditions. The irregular age distribution also appears to reflect the influence of management practices implemented in Norway over past decades. In the following sections, we discuss the key factors driving the observed patterns.

### Age structure patterns and historical management

The oldest sampled birch tree reached 201 years, closely aligning with the maximum reported age of 216 years for *Betula pubescens* in northern Sweden (Hofgaard 1993). However, birch trees over 200 years old were rare or entirely absent across the five biogeographical zones. The middle age group (35–80 years) dominated most regions of the country, although the differences were less pronounced in the northern boreal (NB) and southern boreal (SB) (Fig S1). A similar pattern was noted by Bandekar & Odland (2017), who reported a scarcity of trees older than 77 years in the northernmost birch forests in Norway. Similarly, Aune et al. (2011) found an average tree age of 70.8 ± 28.1 (yr) at tree lines in Northeast Norway and Northwest Russia.

Previous research suggests that a combination of environmental and anthropogenic factors has influenced the age structure of birch forests in Norway. The interplay between topography and strong winds, particularly in the coastal and high-altitude regions of Finnmark, may restrict the establishment and longevity of birch stands (Heikkinen et al. 2002, Bryn et al. 2013). Additionally, human activities have played a critical role in altering stand dynamics. While birch is not a primary target of commercial forestry, selective logging, firewood collection, and land use changes have contributed to the scarcity of older trees, particularly in managed landscapes (Solberg et al. 2005).

These trends align with findings from other regions, where intensive forest management has led to shifts in age class distributions. For instance, Riechelman et al. (2020) observed a decline in old-growth Larix decidua due to harvesting, while Wulder et al. (2009) linked the prevalence of young forests on Vancouver Island to ongoing logging. Similar patterns emerged across Europe following extensive post- war timber exploitation (Gold et al. 2006). In Norway, large-scale afforestation and reforestation efforts were initiated in response to early 20th-century overharvesting, primarily targeting coniferous species but indirectly influencing birch regeneration dynamics (Barth 1916, Landsskogtakseringen 1920, 1933, 1938, 1959, 1968). In 1964, an ambitious national goal was set to double forest production capacity within a century, aiming to increase annual timber growth from 13 to 24 million cubic meters. This growth was achieved across all major tree species, including spruce, pine, and deciduous trees (Landsskogtakseringen 1970). Our findings suggest that the prevalence of the middle age class in Norway’s forests today is a direct result of these interventions. Large-scale planting, afforestation, and improved management practices not only addressed past overharvesting but also shaped the current composition and age distribution of Norway’s forests, reflecting the long-term impact of sustainable management strategies.

Moreover, our results also revealed a relationship between tree age and site index values, with the oldest trees predominantly occurring in areas characterized by lower site index. This association may also reflect, to some extent, the influence of forest management practices on forest development and age distribution. The predominance of younger forests in highly productive sites likely resulted from intensive management strategies aimed at maximizing timber yield and economic returns. In addition, areas with a high site index are often subject to more frequent disturbances or human interventions, including precommercial thinnings and frequent harvesting cycles, which particularly affect birch as an early successional species by limiting its ability to reach advanced. Nonetheless, since the samples for this study were obtained from birch-dominant stands, the observed patterns primarily reflect areas where birch is a significant component of the forest landscape, reducing the influence of management practices targeting other species.

Conversely, less productive sites tend to be managed less intensively, allowing trees to grow for longer periods and potentially reach older ages. Previous research has shown that site productivity, along with tree composition and habitat type, influence growth, lifespan, and turnover rates in forest ecosystems (Stephenson and Mantgem 2005, Di Filippo et al. 2012, Di Filippo et al. 2015). For example, Ziaco et al. (2012) observed that stands at lower elevations, such as Norway’s oceanic west coast, developed faster than those in higher mountain regions, where trees live longer but grow slowly. Our findings align with these observations and those of Di Filippo et al. (2015), highlighting the significant influence of ecological site conditions on tree longevity through their effects on growth rates. The impact of site conditions on growth rates across different age classes will be further explored in detail in the following section.

### Factors influencing age and growth

When comparing trees at the same biological age (e.g., when both an older and a younger tree were 10 years old) (Fig. 4), we found that the newer generation exhibited significantly better growth. This observed radial growth acceleration appears to be particularly pronounced in areas with a high site index, where favorable environmental conditions—such as better soil quality, higher nutrient availability, and milder climatic factors—support enhanced growth.

The observed differences in growth patterns between older and younger birch trees are likely influenced by shifts in environmental conditions over recent decades, including changes in climate and forest structure, such as canopy shape and density (Pretzsch et al. 2014). Similar patterns have been documented in other regions and species. For instance, Johnson and Abrams (2009) reported that younger trees of various species in the United States grew faster at the same biological age compared to older trees over the past 50 to 100 years, suggesting a response to changing environmental factors. In birch specifically, Vanhellemont et al. (2016) highlighted tree age as a critical factor influencing growth in their study in Flanders. Likewise, Hein et al. (2009) demonstrated a strong negative relationship between age and birch growth, with growth rates declining rapidly in trees older than 25 years. This decline may be partly explained by increased competition for resources such as light, which is particularly detrimental for birch, a shade-intolerant pioneer species. The reduced ability of older trees to compete effectively under dense canopy conditions likely exacerbates this growth decline, emphasizing the importance of light availability in birch-dominated stands.

These findings suggest that shifting environmental conditions, including climate change and alterations in forest structure, have played a significant role in shaping birch growth dynamics. Additionally, modern forest management strategies, which often prioritize faster-growing and more productive stands, may reinforce these patterns by favoring younger cohorts in managed forests. As forest practices continue to evolve in response to economic and ecological objectives, understanding the long-term implications of these trends—particularly in the context of climate change—will be essential for sustainable management and conservation efforts.

### The impact of sampling and survivor bias

Recognizing the challenges posed by sampling and survivor bias is critical in our study, as these factors could significantly influence our findings. Survivor bias, in particular, may affect our observations, especially regarding older trees (Bowman et al. 2013, Vanhellemont et al. 2016). In this age class group, fast-growing juvenile trees may be underrepresented in our dataset (Searle and Chen 2018), as many likely died before we could sample them. In contrast, the "young" age class group captures the full range of early growth rates, including both slow and fast-growing individuals. Consequently, faster-growing trees remained in the dataset, potentially increasing the average growth rate observed in younger trees. This could help explain the trends shown in Figure 4, although, as discussed, environmental changes (Pretzsch et al. 2023) might also have contributed to the improved growth of younger trees.

Additionally, our sampling method focused on dominant and co-dominant trees, excluding birch trees with very low growth rates. This selective approach means our results primarily reflect trends in canopy trees, potentially overestimating growth rates compared to the overall population. This may have skewed the relationship between age and growth and affected the interpretation of birch age distribution across the landscape. By excluding stands where birch was not the dominant species (e.g., mixed stands), our analysis might show a biased picture of birch age distribution across diverse forest types.

Recent studies have noted a significant occurrence of missing rings in birch trees in Norway (Harr et al. 2021), often due to environmental stressors such as late frosts and pest outbreaks (Karlsson et al. 2004, Jepsen et al. 2008, Levanič and Eggertsson 2008) as well as other disturbances (Hartl et al. 2019). This introduces the risk of underestimating the ages of older trees, particularly in habitats where ring suppression is more common, such as areas with frequent insect outbreaks. Despite these limitations, the data from the Norwegian National Forest Inventory (NFI), selected purely on statistical criteria (Breidenbach et al. 2020), lack investigator bias and reflect a wide range of ecological conditions across the country. As a result, the dataset provides a comprehensive representation of Norway’s birch population, encompassing trees of varying ages and growth stages, and offers a reliable basis for understanding the age distribution of birch trees across the country. Therefore, we consider this limitation (the potential presence of missing rings) to have minimal impact on the study, given its primary focus on assessing regional patterns in age distribution.

## Acknowledgements

This research was supported by the Research council of Norway (Grant 301922 to the project NORTHERN FOREST. We thank the Norwegian National Forest Inventory (NNFI) field team for their contribution in collecting and processing the tree ring data.

## Author contribution

Marta Vergarechea: Formal analysis, Methodology, Visualization, Writing – original draft. Clara Antón-Fernández: Methodology, Writing -review & editing-, conceptualization. Jane Jepsen: Methodology, Writing -review & editing-, conceptualization. Ole Petter Laksformo Vindstad: Methodology, Writing - review & editing-, conceptualization. Rasmus Astrup: Writing -review & editing-, conceptualization, Funding acquisition.

## Declaration of generative AI and AI-assisted technologies in the writing process

During the preparation of this work the author(s) used ChatGPT in order to revise grammar. After using this tool/service, the authors reviewed and edited the content as needed and take full responsibility for the content of the publication.

## Supplementary materials

**Figure S1:**
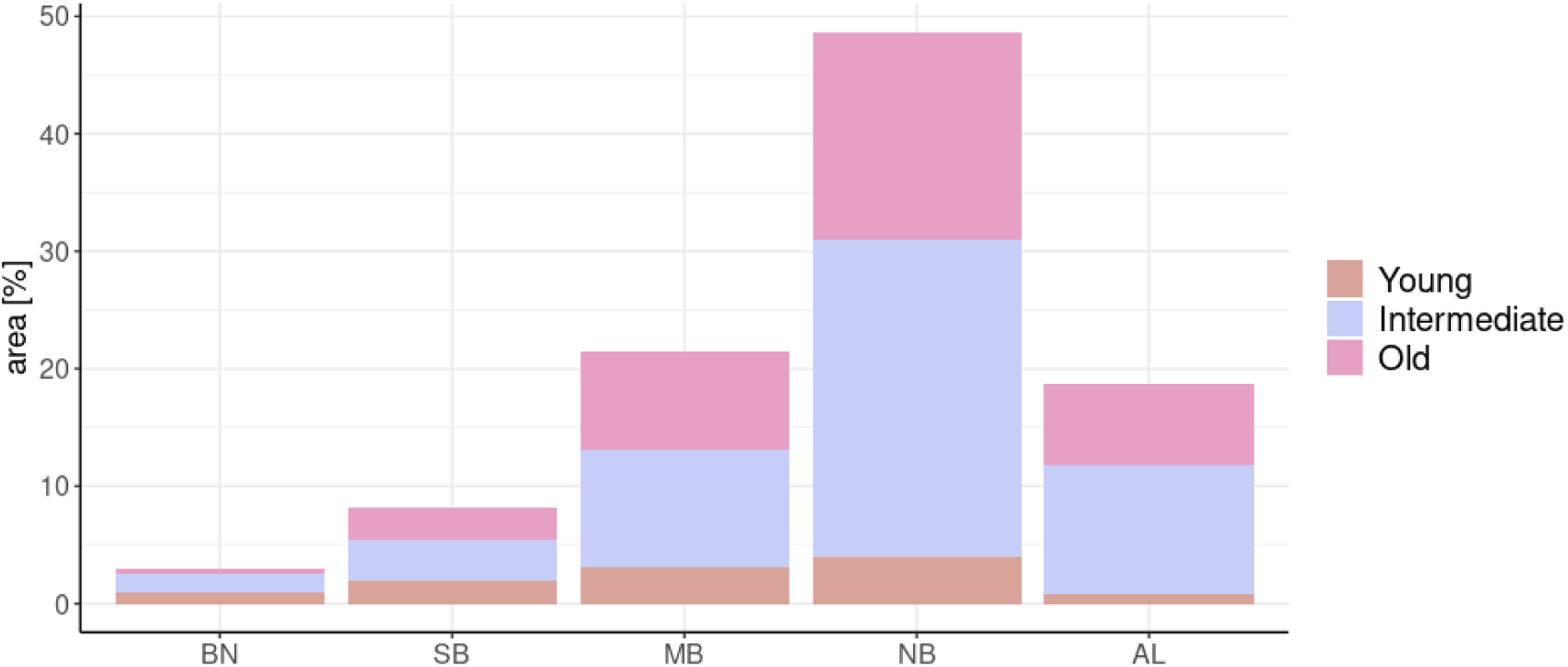
Distribution of birch forests in Norway categorized into three age groups: young (<35 years), old (>80 years), and intermediate- (35-80 years). The data is presented as a percentage of the total birch forest area in Norway. Here BN = boreonemoral, SB = southern boreal, MB = middle boreal, NB = northern boreal and AL = Alpine.

**Figure S2:**
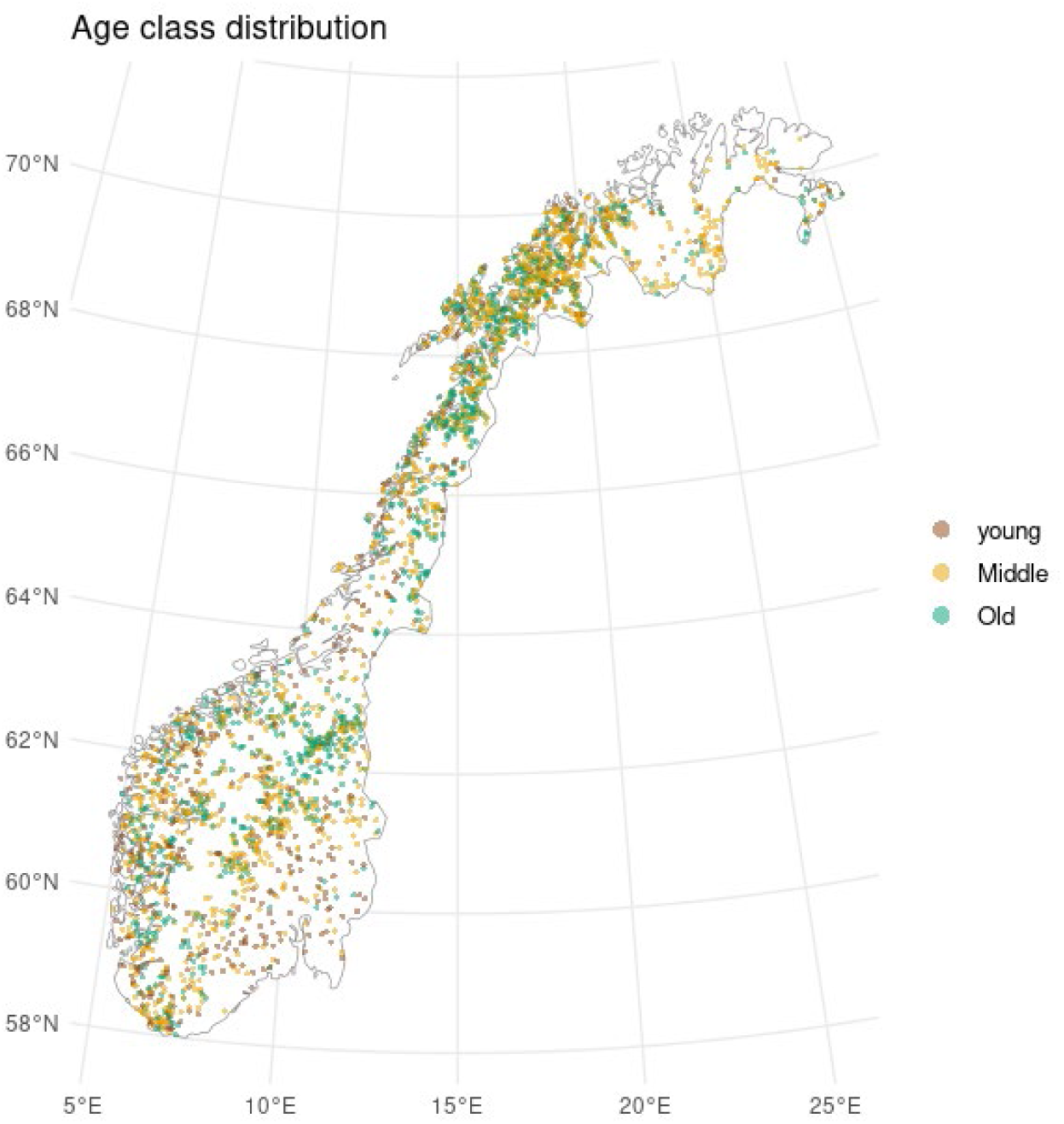
Spatial distribution of birch species age classes across Norway

**Figure S3:**
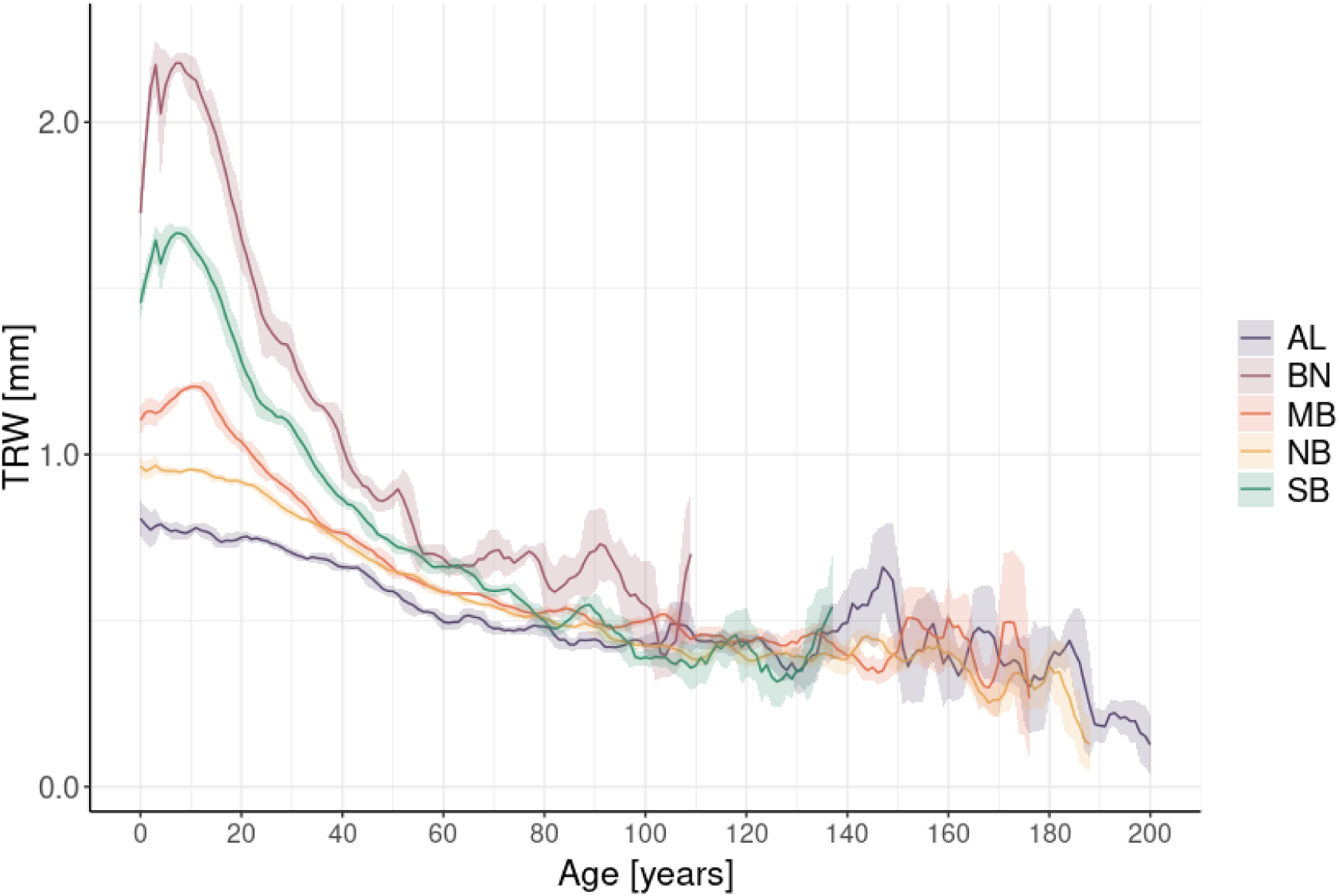
Growth trends for different vegetation (biogeographical) zones. The shaded bands represent mean ± standard error.

**Figure S4:**
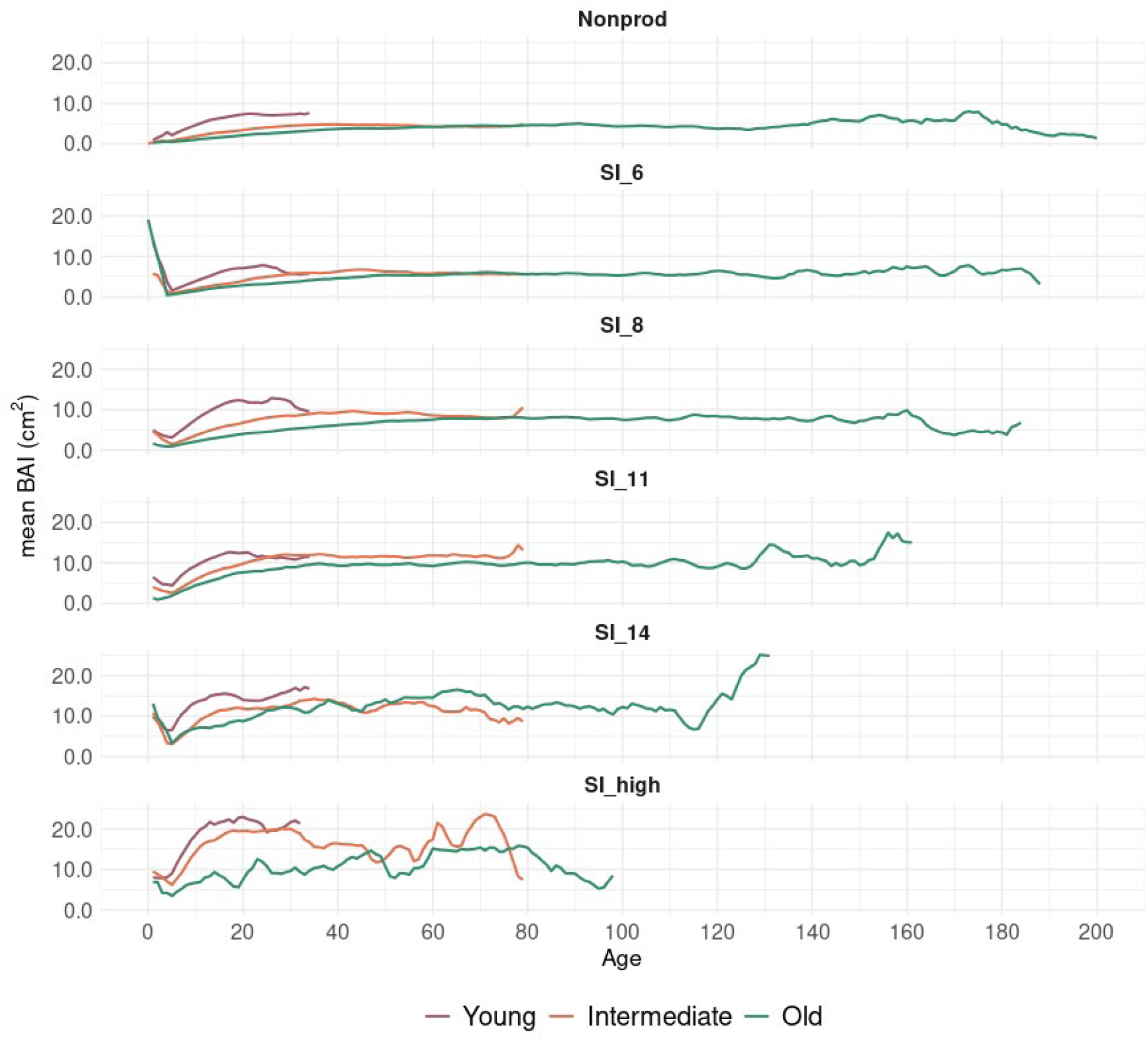
Basal area trends for different site index classifications and young (≤ 50 yrs), intermediate (50-100 yrs) and old age (> 100 yrs) classes.

**Table S1.**
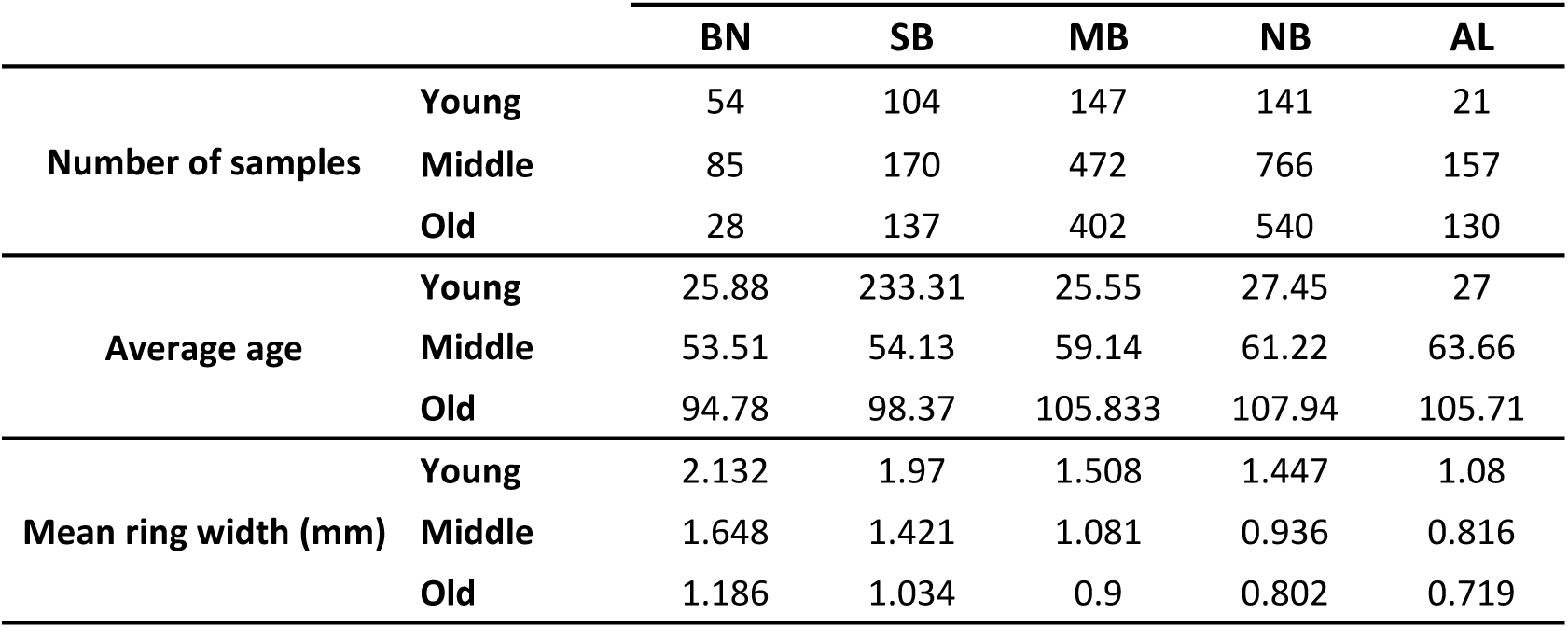
Descriptive summary of the tree ring series.

|  |  | BN | SB | MB | NB | AL |
| --- | --- | --- | --- | --- | --- | --- |
| Number of samples | Young | 54 | 104 | 147 | 141 | 21 |
|  | Middle | 85 | 170 | 472 | 766 | 157 |
|  | Old | 28 | 137 | 402 | 540 | 130 |
| Average age | Young | 25.88 | 233.31 | 25.55 | 27.45 | 27 |
|  | Middle | 53.51 | 54.13 | 59.14 | 61.22 | 63.66 |
|  | Old | 94.78 | 98.37 | 105.833 | 107.94 | 105.71 |
| Mean ring width (mm) | Young | 2.132 | 1.97 | 1.508 | 1.447 | 1.08 |
|  | Middle | 1.648 | 1.421 | 1.081 | 0.936 | 0.816 |
|  | Old | 1.186 | 1.034 | 0.9 | 0.802 | 0.719 |

